# A single-antigen, multi-epitope subunit vaccine candidate against lumpy skin disease virus designed by conservation-guided reverse vaccinology

**DOI:** 10.64898/2026.08.16.745068

**Authors:** Bidhan Koirala

## Abstract

Lumpy skin disease virus causes devastating economic losses in cattle, and current live attenuated vaccines carry risks of reversion and cannot differentiate infected from vaccinated animals. We applied conservation-guided reverse vaccinology (vaccine design from genomic sequence data) to identify conserved epitopes (immune-recognized protein fragments) and design a single-antigen, multi-epitope subunit vaccine candidate. A nine-taxon phylogenetic supermatrix of three candidate antigens was built, and BLA-restricted T cell and linear B cell epitopes were predicted. Conservation was quantified via Shannon entropy and Fisher’s exact tests; structural disorder via AlphaFold2 and IUPred2A, followed by codon optimization and in silico cloning. All six selected epitopes mapped to the ankyrin locus. The 134 epitope columns showed significantly higher constraint than 2,376 background columns (mean entropy 0.008 vs. 0.243 bits; odds ratio 29.74). The 87-aa, 9.31 kDa construct was predominantly disordered and cloned in silico into pET28a. This work is entirely computational; wet-lab validation of immunogenicity is required before translational claims.

## 1. INTRODUCTION

Lumpy skin disease (LSD), caused by the Lumpy Skin Disease Virus (LSDV; family Poxviridae, genus Capripoxvirus), has emerged as one of the most economically consequential transboundary diseases of cattle over the past two decades. First described in Zambia in 1929 and long confined to sub-Saharan Africa, LSDV has undergone a dramatic geographic expansion since 2012, sweeping through the Middle East, Turkey, the Balkans, the Caucasus, and—most recently—the Indian subcontinent and Southeast Asia (Tuppurainen et al., 2011; Klement, 2015; Bhanuprakash et al., 2013; Sudhakar et al., 2023). Clinical disease is characterised by fever, cutaneous nodules, lymphadenopathy and reduced milk production, with mortality occurring in a proportion of affected animals. Even when mortality is limited, morbidity and production losses can result in substantial economic effects. (Babiuk et al., 2008). The Food and Agriculture Organization estimates that the 2019–2022 Indian epizootic alone affected over two million cattle and caused losses exceeding 18 billion INR, underscoring the urgency of effective prophylactic strategies (FAO, 2022; WOAH, 2023).

Control of LSD still rests largely on live-attenuated vaccines, most notably the Neethling strain, which, although protective, carries well-documented drawbacks: residual virulence, reversion risk upon serial passage, cold-chain dependency, and an inability to differentiate infected from vaccinated animals (DIVA) (Abutarbush, 2014; Russell et al., 2018). In many newly affected regions, heterologous sheeppox- or goatpox-based vaccines are deployed out of necessity, yet cross-protection data remain inconsistent and regulatory approval is patchy (Tuppurainen and Lyra, 2011). These limitations have renewed interest in rational, sequence-driven vaccine design that circumvents whole-pathogen manipulation entirely.

Reverse vaccinology and immunoinformatics provide an in silico approach for identifying conserved antigens and candidate immune-reactive peptides. Genome-based analyses can be used to identify conserved antigenic regions, predict MHC-binding peptides, and combine selected epitopes into a candidate chimeric immunogen before experimental evaluation. (Sette and Rappuoli, 2010; De Groot et al., 2008). For LSDV, three targets have attracted particular attention: the G-protein-coupled receptor (GPCR), involved in host-cell tropism and immune evasion; the extracellular enveloped virion (EEV) glycoprotein, a key determinant of cell-to-cell spread; and an Ankyrin-repeat protein implicated in modulating innate antiviral responses (Tulman et al., 2001; Gao et al., 2017). Because LSDV exhibits relatively low nucleotide diversity across its genome, these antigens are attractive candidates for a pan-strain subunit vaccine, provided that epitope-level conservation can be demonstrated quantitatively rather than assumed.

A further practical consideration concerns the bovine immune context. Cattle MHC class I molecules, designated bovine lymphocyte antigen (BoLA), are highly polymorphic, and any credible T-cell vaccine must present epitopes across a representative panel of BoLA alleles to ensure population-level coverage (Davies et al., 2012; Sanz-Parra et al., 1999). Also, structural assessment can provide additional information on the predicted organization and accessibility of epitopes within a multi-epitope construct. AlphaFold2 (Jumper et al., 2021), including its ColabFold implementation (Mirdita et al., 2022), provides an additional computational approach for assessing the predicted structural properties of vaccine constructs. Codon optimisation for Escherichia coli and in silico cloning into an expression vector such as pET-28a(+) can also be used to evaluate the feasibility of subsequent recombinant production (Nezafat et al., 2016).

The objective of this study was to identify conserved LSDV epitopes and use them to design a multi-epitope subunit vaccine candidate through a combined phylogenetic, immunoinformatics and structural analysis. To that end, the work was organised around five specific aims: (i) to construct a nine-strain phylogenetic supermatrix and identify highly conserved target proteins (GPCR, EEV, and Ankyrin) across geographically diverse LSDV isolates; (ii) to predict BoLA-restricted cytotoxic T-lymphocyte epitopes and linear B-cell epitopes and validate their evolutionary conservation through Shannon entropy analysis; (iii) to assemble the most conserved and promiscuous epitopes into a single chimeric protein using GPGPG linkers and evaluate its physicochemical and safety profile; (iv) to predict the three-dimensional structure of the final construct via AlphaFold2, confirming solvent exposure of B-cell epitopes and overall scaffold integrity; and (v) to codon-optimise the construct for E. coli expression and perform in silico cloning into the pET-28a(+) vector to demonstrate readiness for downstream wet-lab manufacturing.

## 2. MATERIALS AND METHODS

### 2.1. Sequence data and supermatrix construction

Complete coding sequences of three LSDV antigens (GPCR chemokine receptor homologue, extracellular enveloped virus (EEV) homologue and ankyrin repeat protein) were retrieved from GenBank for seven LSDV strains (NC_003027.1, KX764645.1, OQ588787.1, OR393176.1, OR393177.1, OR863389.1 and PQ616985.1) and two outgroup capripoxviruses (sheeppox virus NC_004002.1 and goatpox virus NC_004003.1). Accessions OR797612.1, PP979138.1 and PQ472735.1 were excluded because one or more loci were missing; taxa lacking complete loci cannot be placed reliably in a concatenated analysis. Each locus was aligned with MUSCLE (Edgar, 2004) in MEGA 12 (MEGA, 2025) and the three alignments were concatenated in the order GPCR, EEV, ankyrin with a custom Python script to produce a 2,526 bp supermatrix (GPCR 1,245 bp; EEV 645 bp; ankyrin 636 bp; Supplementary File S1).

### 2.2. Phylogenetic analysis

A maximum-likelihood tree was inferred from the supermatrix in MEGA 12 (MEGA, 2025) with 1,000 nonparametric bootstrap replicates; the substitution model was selected by the Bayesian information criterion within MEGA. The tree was rooted with the sheeppox and goatpox outgroups and annotated by country of origin in the Interactive Tree Of Life (iTOL) (Letunic and Bork, 2021).

### 2.3. Epitope prediction

Nine-mer T-cell epitopes were predicted with NetMHCpan 4.1 against BoLA class I alleles (Reynisson et al., 2020), and peptides meeting the recommended binding ranks were retained. Linear B-cell epitopes were identified with a consensus pipeline combining hydrophilicity, surface accessibility and antigenicity scores. Candidate epitopes were then filtered for evolutionary conservation as described in section 2.4.

### 2.4. Epitope conservation analysis

Per-column Shannon entropy (Shannon, 1948) was computed across the nine-taxon supermatrix with a custom Python script (NumPy (Harris et al., 2020), SciPy (Virtanen et al., 2020)). Epitope coordinates were mapped to supermatrix columns by gap-aware, codon-aligned translation of the reference sequence (NC_003027.1), and overlapping intervals were merged to define the set of unique epitope columns; all remaining columns served as the background. Sixteen alignment columns consisting entirely of gap characters were removed before statistical analysis. The analysed supermatrix therefore contained 2,510 columns: 134 epitope columns and 2,376 background columns. Gap characters in partially gapped columns were treated as missing data. A column was classified as invariant when all non-gap nucleotides were identical. Entropy distributions of epitope and background columns were compared with a two-sided Mann-Whitney U test (Mann and Whitney, 1947), and enrichment of fully invariant columns (entropy = 0) was tested with Fisher’s exact test (Fisher, 1922). Invariance was additionally scored per epitope as the percentage of entropy-zero columns within its span. All tests were two-sided, with significance set at p < 0.05.

### 2.5. Vaccine construct design and physicochemical analysis

The six epitopes were assembled in the order T1, T2, T3, T4, B1, B2, separated by GPGPG linkers, with a 6×His tag fused at the C-terminus, yielding an 87-residue construct. Molecular weight, theoretical pI and instability index were computed for the mature 87-residue construct with the ProtParam algorithm (Gasteiger et al., 2005).

### 2.6. Codon optimization and in silico cloning

The construct was reverse-translated and codon-optimized for E. coli with a custom Python script (Biopython (Cock et al., 2009)) applying the E. coli K-12 codon usage table (Nakamura et al., 2000) and selecting the most frequent synonymous codon for each amino acid. The 261-bp coding sequence was designed without an initiator ATG; the ATG of the 5’ NdeI site (CATATG) serves as the start codon, and a TAA stop codon followed by the 3’ XhoI site (CTCGAG) completes the 276-bp insert for directional cloning into pET-28a(+) under T7 RNA polymerase control (Studier and Moffatt, 1986). Restriction sites, including Type IIS cut positions, were determined with the Biopython Bio.Restriction module (REBASE database (Roberts et al., 2015)), and the NdeI/XhoI/BfuAI diagnostic digest was computed from the exact cut coordinates.

### 2.7. Structural and disorder assessment

The construct was modeled with AlphaFold2 (Jumper et al., 2021) as implemented in ColabFold (Mirdita et al., 2022) under default multiple-sequence-alignment and template settings. Per-residue pLDDT values and the predicted aligned error were extracted from the top-ranked model, which was visualized in UCSF ChimeraX (Pettersen et al., 2021). Intrinsic disorder was predicted independently with IUPred2A (long mode) using a disorder probability cutoff of 0.5 (Meszaros et al., 2018).

### 2.8. Software and statistics

Custom steps were implemented in Python 3.14 (NumPy (Harris et al., 2020), SciPy (Virtanen et al., 2020), Biopython (Cock et al., 2009), matplotlib (Hunter, 2007)). Statistical procedures are described in section 2.4. All sequence accessions and supplementary files are listed in the Data Availability statement.

## 3. RESULTS

### 3.1. The three candidate antigens show high sequence conservation among the analyzed LSDV strains

The GPCR, EEV and ankyrin loci were aligned separately with MUSCLE in MEGA 12 and concatenated into a 2,526 bp alignment (GPCR 1,245 bp; EEV 645 bp; ankyrin 636 bp) for seven LSDV strains plus sheeppox and goatpox as outgroups. Three GenBank sequences were omitted because they were missing one or more loci (OR797612.1, PP979138.1, PQ472735.1). Taxa lacking one or more of the three loci were excluded from the concatenated analysis because missing loci would result in incomplete representation of the supermatrix. In the maximum likelihood tree (1,000 bootstrap replicates; Fig. 1), the outgroups separated from the LSDV strains with bootstrap 100 and the LSDV strains formed a single clade. Within the LSDV clade, the Kenyan strain KX764645.1 occupied a basal position (74), followed by one Indian strain (OR393176.1; 81) and the South African vaccine strain Neethling (NC_003027.1; 51); the remaining Indian strains and the Pakistani strain grouped terminally (98, 90). Support for several deeper nodes was low (51 to 81), limiting resolution of relationships within the clade.

**Figure 1.**
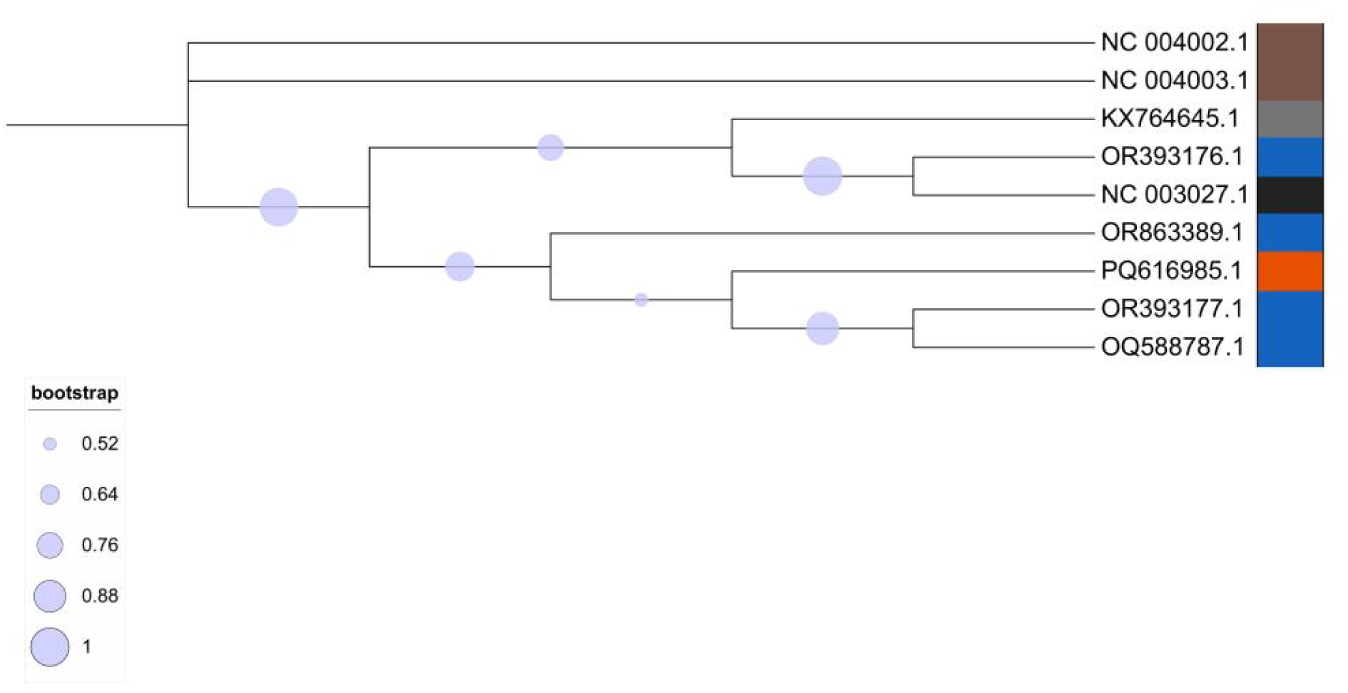
Maximum-likelihood phylogenetic tree of LSDV strains. The tree was inferred from a concatenated supermatrix (GPCR, EEV, and ankyrin genes; 2,526 bp) and rooted with sheeppox/goatpox outgroups. Nodes are labeled with light-blue circles proportional to bootstrap support (1,000 replicates). Taxon names are colored according to species classification.

### 3.2. The selected epitopes are evolutionarily constrained

Four T-cell epitopes (EEDENGKNL, DENGKNLLH, YETPLFSCV and KIFDYLTSL; Table 1) were retained from the NetMHCpan predictions against BoLA class I alleles. Two linear B-cell epitopes (NFLQKDFDNN and LFHSDKKIFD; Table 2) were selected from the consensus hydrophilicity, accessibility and antigenicity pipeline. All six epitopes mapped within the ankyrin locus, spanning supermatrix columns 2043 to 2435. The GPCR and EEV loci contributed no selected epitopes and served as conservation controls in the phylogenetic analysis.

**Table 1.** Predicted BoLA-restricted T-cell epitopes selected for the vaccine construct (invariance computed across all nine taxa).

| Epitope | Source antigen | Construct position | Supermatrix columns | Invariant (%) |
| --- | --- | --- | --- | --- |
| EEDENGKNL | ankyrin | 1-9 | 2244-2270 | 100 |
| DENGKNLLH | ankyrin | 15-23 | 2250-2276 | 96 |
| YETPLFSCV | ankyrin | 29-37 | 2043-2069 | 93 |
| KIFDYLTSL | ankyrin | 43-51 | 2409-2435 | 100 |

**Table 2.** Predicted linear B-cell epitopes selected for the vaccine construct (invariance computed across all nine taxa).

| Epitope | Source antigen | Construct position | Supermatrix columns | Invariant (%) |
| --- | --- | --- | --- | --- |
| NFLQKDFDNN | ankyrin | 57-66 | 2341-2370 | 100 |
| LFHSDKKIFD | ankyrin | 72-81 | 2392-2421 | 97 |

The 134 unique nucleotide columns encoding the six selected epitopes (overlapping positions counted once) carried almost no variation. Because the highly conserved LSDV background produced a median Shannon entropy of 0.000 in both epitope and background columns, conservation was additionally evaluated by comparing mean Shannon entropy and the proportion of fully invariant positions. Mean Shannon entropy was 0.008 bits in the 134 epitope columns and 0.243 bits in the 2,376 background columns (Mann-Whitney U = 111,422.0, p = 2.01 × 10⁻¹³; Fig. 2). Fully invariant positions comprised 132 of 134 epitope columns (99%) and 1,638 of 2,376 background columns (69%) (Fisher’s exact test, OR = 29.74, p = 2.37 × 10⁻¹⁸). Evaluated individually, the T-cell epitopes were 93 to 100% invariant, and the B-cell epitopes were 97 to 100% invariant across the nine taxa. The selected epitopes therefore reside under strict evolutionary constraint, reducing the likelihood of sequence-based immune escape.

**Figure 2.**
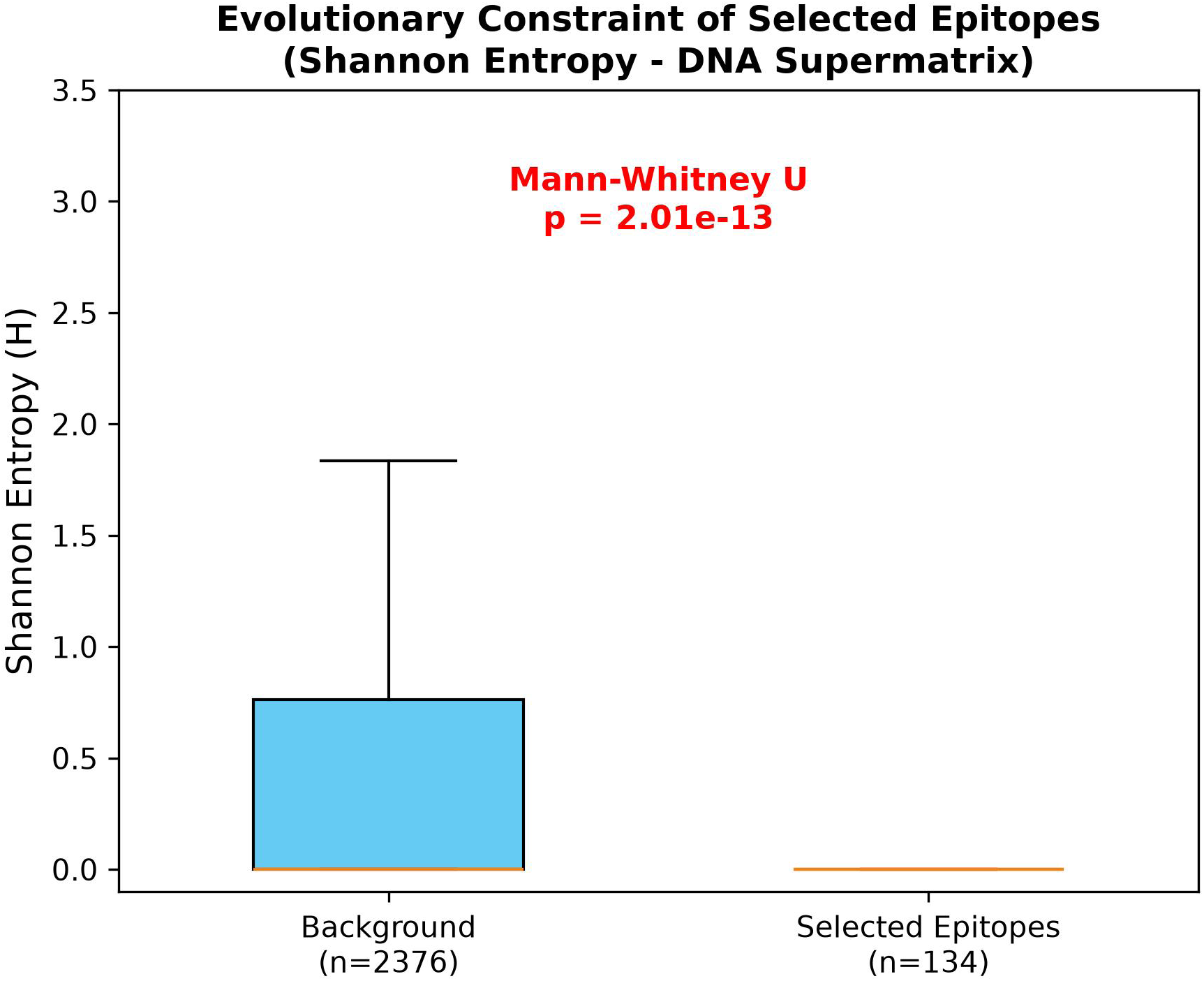
Evolutionary constraint of the selected epitopes. Per-column Shannon entropy (H, bits) across the nine-taxon DNA supermatrix for the 134 epitope columns versus 2,376 background columns (two-sided Mann-Whitney U, p = 2.01 × 10⁻¹³; Fisher’s exact OR = 29.74, p = 2.37 × 10⁻¹⁸).

### 3.3. Construct design

The final 87-amino acid construct assembles the four selected T-cell epitopes (residues 1-9, 15-23, 29-37, 43-51) and two B-cell epitopes (residues 57-66, 72-81), separated by GPGPG linkers, with a 6×His tag at the C-terminus (residues 82-87) (Fig. 3). The construct has an instability index of 8.11, a molecular weight of 9.31 kDa and a theoretical pI of 5.63 (computed for the 87-aa mature construct). No linker separates the 6×His tag from the C-terminal B-cell epitope; possible effects on LFHSDKKIFD accessibility will be assessed, and a GGSGGS spacer may be introduced if required. The construct is small (9.31 kDa); fusion partners such as SUMO or MBP may be evaluated in subsequent expression experiments to improve stability and solubility. The current design prioritizes BoLA class I restricted cytotoxic T cell epitopes and linear B cell epitopes; CD4 helper epitopes may be added in future iterations.

**Figure 3.**
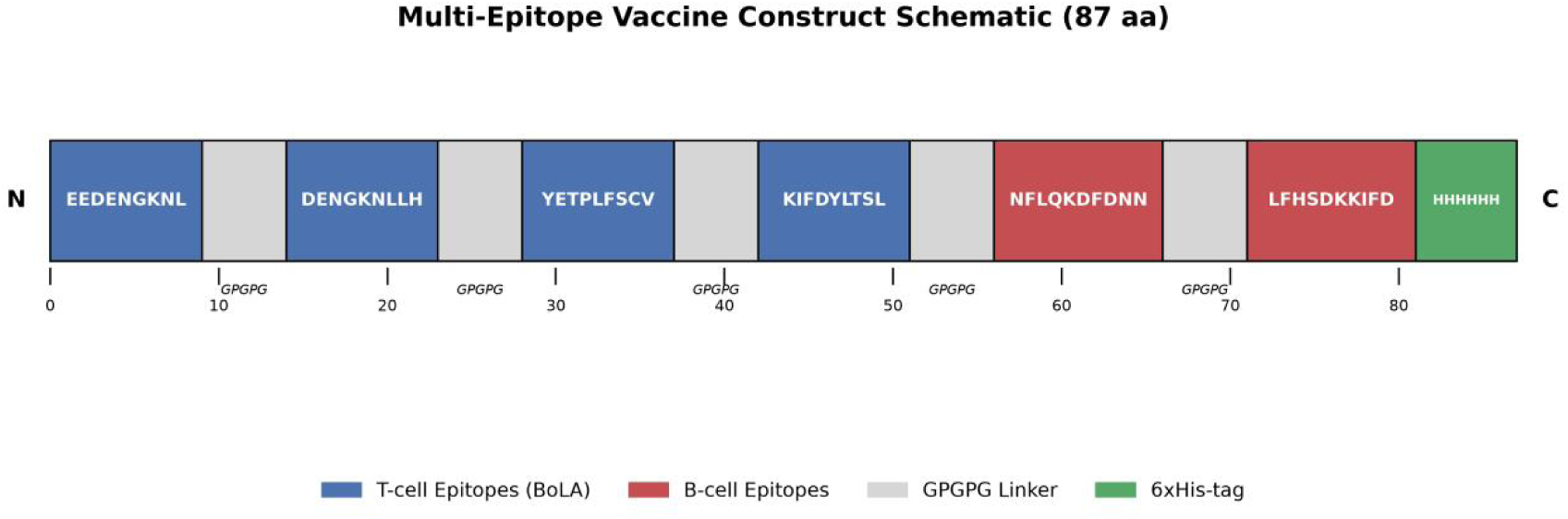
Schematic representation of the multi-epitope vaccine construct. The 87-amino acid construct comprises four conserved T-cell epitopes (blue) and two B-cell epitopes (red), adjoined by flexible GPGPG linkers (grey). A 6×His tag (green) was fused at the C-terminus to facilitate downstream protein purification.

### 3.4. Structural and disorder assessment

AlphaFold2 returned a predominantly flexible model, with residues in the 50 to 70 and <50 pLDDT bands (Fig. 4B,C) and low long-range coordinate correlation in the PAE (Fig. 4D). A single alpha-helix overlaps the T4 epitope (residues 43 to 51; Fig. 4A). Both B-cell epitopes mapped to solvent-exposed segments (Fig. 4E,F). IUPred2A predicted 73 of 87 residues (84%) as intrinsically disordered using a disorder probability cutoff of 0.5, overlapping the low-confidence regions of the AlphaFold2 model. Because most of the model has low pLDDT, solvent exposure of the B-cell epitopes was inferred from predicted disorder and sequence accessibility rather than from high-confidence folded coordinates. Molecular dynamics simulations and experimental circular dichroism are needed to confirm the flexible scaffold.

**Figure 4.**
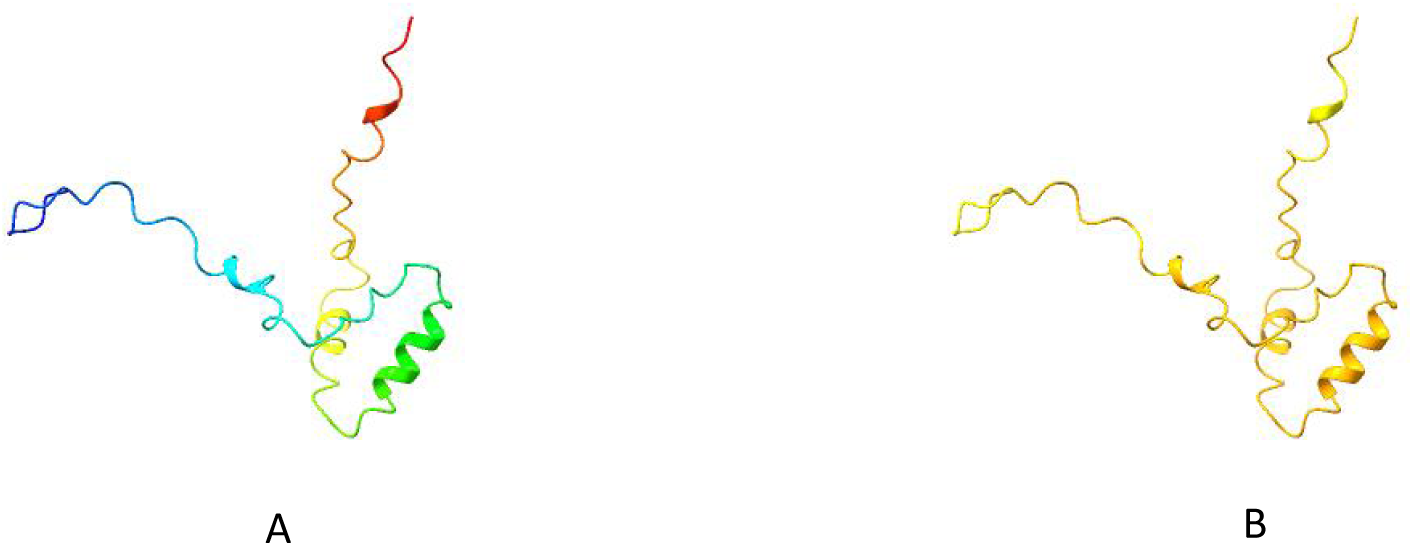
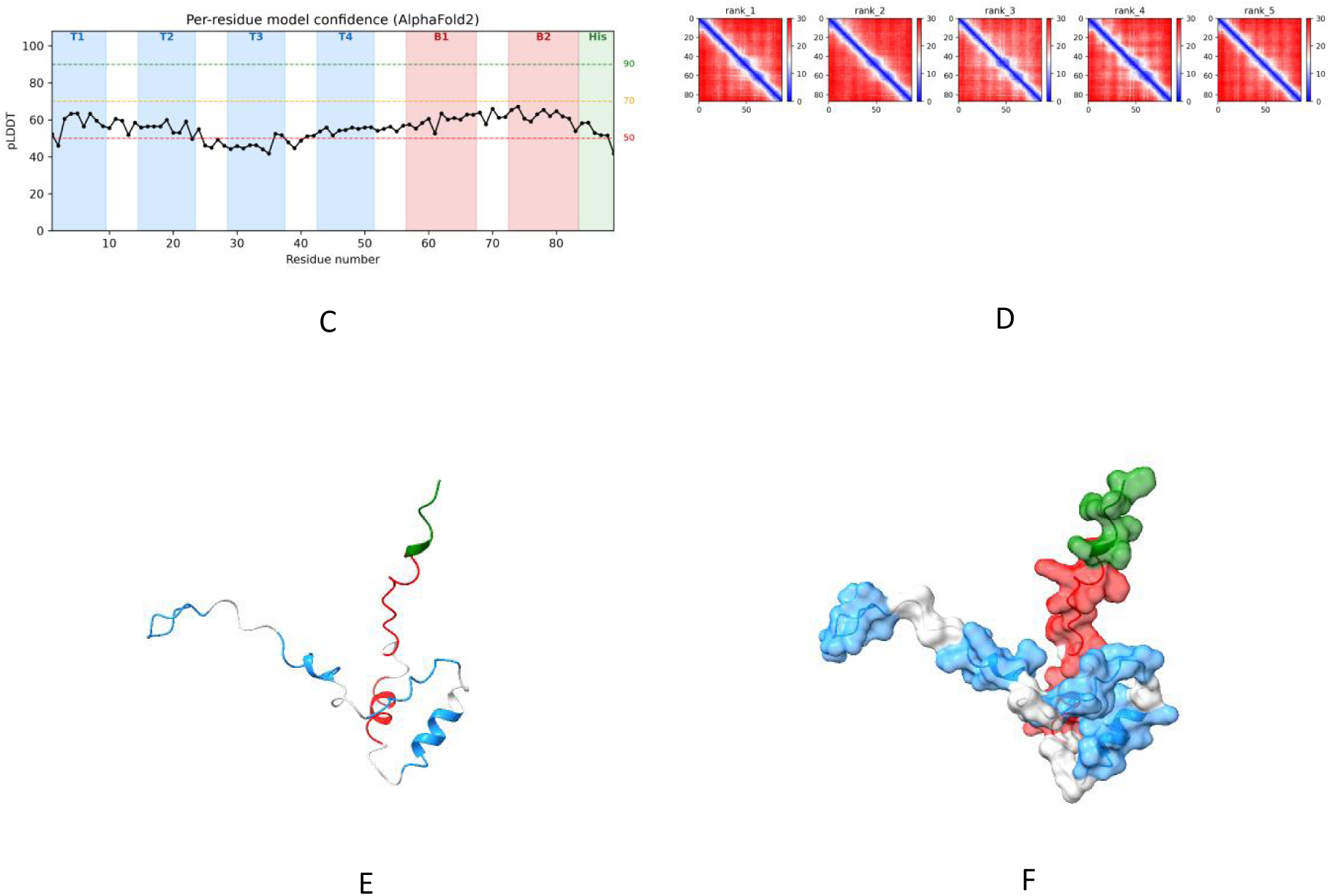
AlphaFold2-predicted structure of the LSDV vaccine candidate. (A) Rainbow cartoon from the N terminus (blue) to the C terminus (red); the single alpha helix (residues 43 to 51) corresponds to the green segment of the gradient. (B) Model colored by per-residue pLDDT (yellow 50 to 70; orange < 50). (C) Per-residue pLDDT profile. (D) PAE heatmap. (E) Epitope mapping on the cartoon model (T-cell, blue; B-cell, red). (F) Surface representation of the same mapping.

### 3.5. In silico cloning and expression strategy

The 87-amino acid construct was reverse-translated and codon-optimized using the Escherichia coli K-12 codon usage table for expression in E. coli strains such as BL21(DE3), yielding a 261-bp coding sequence beginning at the glutamate codon GAA. The 261-bp coding sequence excludes the initiator ATG; the initiator ATG is contributed by the NdeI site. The sequence was flanked by NdeI (CATATG) and XhoI (CTCGAG) sites, and a TAA stop codon was placed downstream of the ORF. The final insert is 276 bp (GC 56.70%). For diagnostic screening, NdeI, XhoI and BfuAI were used together; this releases the vector backbone and two insert fragments of 72 bp and 194 bp (Fig. 5). These fragment lengths reflect the exact enzyme cut coordinates and exclude the terminal recognition site overhangs; therefore, the sum of the resolved insert fragments differs from the annotated 276 bp insert length. The translated product includes an initiator methionine supplied by the NdeI ATG, followed by the 87 residue mature design. The 87-residue construct is the design antigen; because of its small size (9.31 kDa), experimental production will require a fusion partner such as SUMO or MBP.

**Figure 5.**
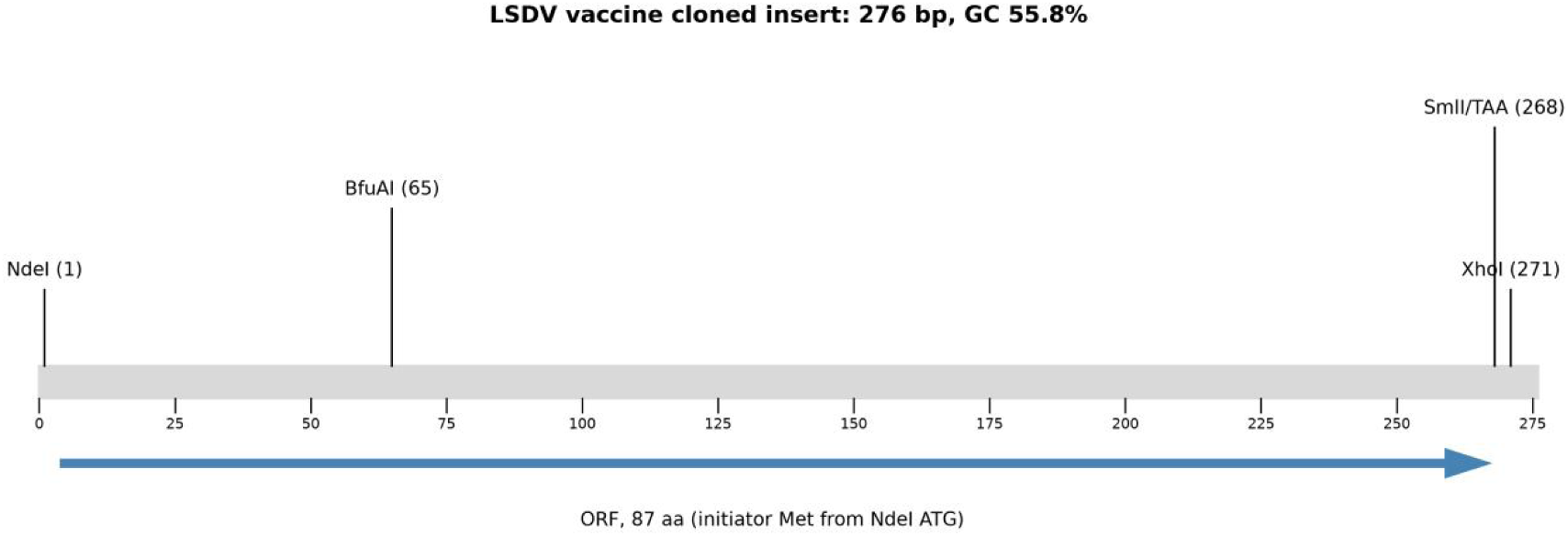
Cloning map of the 276-bp insert (GC 56.70%). NdeI (1) and XhoI (271) flank the insert for directional cloning into pET-28a(+); the ORF (nt 4 to 267) is shown as an arrow. The internal BfuAI site (65) and the TAA stop codon (268 to 270) are indicated.

## 4. DISCUSSION

In this study, conserved regions of three LSDV antigens were screened at the epitope level, and selected epitopes were assembled into a multi-epitope construct that was subsequently evaluated using structural prediction and in silico cloning. Of the three antigens screened, the ankyrin repeat protein supplied all six selected epitopes, and the final 87-residue construct assembled them with GPGPG linkers and a C-terminal 6×His tag.

The phylogenetic analysis placed the seven LSDV strains in a single clade with limited resolution at deeper nodes. The relatively low bootstrap support at several internal nodes limits the resolution of relationships among the LSDV strains. Nevertheless, the overall clustering of the analysed strains and the high sequence conservation observed across the three loci support their suitability for further epitope-level analysis (Tuppurainen et al., 2011). The entropy and invariance analyses showed that the selected epitope regions contained substantially less sequence variation than the remaining alignment. The high conservation of these regions is relevant to vaccine design because sequence variation within conserved epitopes may be less permissive than variation in more variable regions. However, the present analysis does not directly assess the potential for immune escape (Sprygin et al., 2019).

All selected epitopes mapped to the ankyrin locus. Ankyrin repeat proteins are immunogenic in LSDV-infected cattle (Gao et al., 2017) and are annotated in the Neethling genome as host-range and immune-modulatory genes (Tulman et al., 2001), supporting the biological relevance of the selected regions. The GPCR and EEV loci, although conserved, did not supply epitopes passing the prediction and conservation filters and therefore served as internal conservation controls. Because all six selected epitopes originated from the ankyrin locus, the present construct represents a single-antigen multi-epitope design. Additional epitopes from GPCR or EEV could be evaluated in future designs if broader antigenic representation is required.

The AlphaFold2 model was predominantly low-confidence, and IUPred2A predicted 84% of residues as disordered. The high predicted disorder is consistent with the short, linker-rich composition of the construct. GPGPG linkers are commonly used to provide flexibility between adjacent epitopes (Chen et al., 2013), although the present analysis cannot establish whether this flexibility improves epitope processing or B-cell recognition. The low-confidence regions predicted by AlphaFold2 overlapped substantially with regions assigned high disorder probability by IUPred2A, providing consistent computational evidence for a flexible construct. The predicted exposure of the B-cell epitopes should therefore be interpreted cautiously because most of the surrounding structural model had low confidence. Molecular dynamics and circular dichroism are required to characterize the conformational ensemble.

The proposed cloning strategy uses NdeI and XhoI sites for directional insertion into pET-28a(+), while the predicted BfuAI digest provides an additional in silico check of the construct (Studier et al., 1990; Roberts et al., 2015). Two limitations of the current construct are apparent. The mature construct is only 9.31 kDa, a size at which E. coli proteases degrade unprotected peptides; expression will therefore require a fusion partner such as SUMO or MBP. In addition, the construct contains only BoLA class I restricted and linear B-cell epitopes; CD4 helper epitopes, which support memory and antibody maturation, are absent and should be incorporated in future designs.

The work is entirely computational. No expression, immunoassay or challenge data were generated, and the BoLA panel, although representative, does not cover the full diversity of cattle MHC (Davies et al., 2012). Because the proposed vaccine contains selected LSDV epitopes rather than a complete infectious virus, it could potentially be developed within a DIVA-compatible strategy. However, DIVA performance would require experimental validation and an appropriate diagnostic assay.

## 5. CONCLUSION

This study used a combined phylogenetic and immunoinformatics approach to identify conserved LSDV epitopes for multi-epitope vaccine design. Six epitopes from the ankyrin locus were selected on BoLA class I binding predictions and hydrophilicity, accessibility and antigenicity consensus, and showed significantly lower Shannon entropy and higher invariance than background alignment columns.

The resulting 87-amino acid construct contained four predicted T-cell epitopes and two linear B-cell epitopes separated by GPGPG linkers. Structural prediction indicated a predominantly low-confidence and disordered model, with the selected B-cell epitopes located in predicted accessible regions.

The work is entirely computational: no wet-lab expression, immune-cell assays, animal challenge or molecular dynamics simulations were performed, and the construct has not been tested for immunogenicity, expression stability or protective efficacy. AlphaFold2 predictions for disordered constructs carry inherent uncertainty, and the small construct will require a fusion partner for reliable expression.

Next steps include molecular dynamics simulation, BoLA and pattern-recognition receptor docking, construction of a fusion-tagged expression cassette, recombinant protein production, in vitro immune stimulation assays and, ultimately, in vivo evaluation in cattle.

These findings identify a conserved multi-epitope construct for further investigation, but its expression, immunogenicity, stability and protective efficacy remain to be determined experimentally.

